# Top-down effects in motor generalization

**DOI:** 10.1101/2021.02.09.430542

**Authors:** Eugene Poh, Naser Al-Fawakhiri, Rachel Tam, Jordan A. Taylor, Samuel D. McDougle

**Author notes:** These authors contributed equally to this work (Jordan A. Taylor, Samuel D. McDougle). Correspondence: Eugene Poh, **Email:**. **Author Contributions:** E.P., J.A.T and S.D.M. designed research; E.P., N.A., and R.T., performed research. E.P., N.A., R.T., J.A.T., and S.D.M. analyzed data; E.P., J.A.T., and S.D.M. wrote the paper.

## Abstract

To generate adaptive movements we must generalize what we have previously learned to novel situations. The generalization of adapted movements has typically been framed as a consequence of neural tuning functions that overlap for similar movement kinematics - what might be considered bottom-up generalization. However, as is true in many domains of human behavior, generalization can also be framed as the result of deliberate decisions about how to act (top-down generalization). Here we attempt to broaden the scope of theories about motor generalization, hypothesizing that part of the typical motor generalization function can be characterized as a consequence of top-down decisions concerning the subjective similarity of different movement contexts. We tested this proposal by having participants make explicit similarity ratings over both traditional kinematic contextual dimensions (movement direction) and more abstract contextual dimensions (target shape), and perform a visuomotor adaptation generalization task where trials varied over those dimensions. Across five experiments, we measured the relationship between subjective similarity ratings and motor generalization. In some cases this link was rather strong, though it was determined by both task-relevance and explicit instruction. These results support a broadening of the descriptive framework used to understand the generalization of motor behaviors and support a more careful deployment of instructions in generalization studies.

**Significance Statement:** Generalization describes the transfer of knowledge from one context to another, and is typically thought to result from a higher-order inference process. However, in the motor adaptation domain, generalization is thought to arise from neural representations tuned to low-level kinematics. To bridge these differing views, we measured peoples’ subjective similarity judgements of different task contexts during sensorimotor adaptation. We found that motor generalization was closely linked to participant’s subjective judgements, and that explicit instructions about the consequential dimension of different contexts further shaped generalization. These findings emphasize that in addition to low level kinematic considerations, top-down inferences about which action to take in a given context should be considered as another key component of motor generalization.

## Introduction

Adaptive motor behavior is not just about executing movements precisely, it is also about selecting them intelligently. For example, in billiards one must select the best approach for a shot (follow or draw?) and then apply a precise amount of force in the stroke. In novel situations, agents must rely on prior experience to select and execute movements effectively. One approach to generalization is to make inferences about how one should move in a new context (1). Indeed, inference is thought to be the brain’s solution for generalizing learned behaviors across diverse environmental contexts (2–4).

In the motor learning domain, however, theories of generalization often focus on physical kinematic considerations rather than cognitive processes. To illustrate, consider the typical generalization experiment in a motor learning study, in which an individual makes center-out reaches in a radial workspace: When a particular movement – say a reach directed at a 90° target – is repeatedly paired with a sensorimotor perturbation, such as a 30° clockwise rotation of a visual feedback signal, the individual will incrementally adapt their movements, eventually directing their reaches toward 120° to restore performance. If the learner is then presented with a novel target – say at 135° – their movement kinematics will still show signatures of adaptation. That is, their reaches might be directed toward ~145°, reflecting partial adaptation even though they never experienced a perturbation at that location. This behavior is presumed to reflect representational principles that are “baked into” the motor system, where neural populations parametrically encode kinematics or muscle dynamics (5–10). In this view, generalization of an adapted sensorimotor map is explained by, for instance, partially overlapping neural responses for similar movements (8, 11).

Experimental investigations of motor generalization across various kinematic variables (e.g. movement direction, distance, limb positions, velocity, etc.) have, for the most part, shown an orderly gradient of decay with increasing differences between the original training stimulus and the novel test stimulus (6, 8–10, 12–20). However, beyond this general characteristic, the specific pattern of the generalization function tends to differ depending on the specific task or condition. That is, across a wide range of motor adaptation tasks (e.g. visuomotor perturbations such as visuomotor rotations, gain changes, translations, prismatic shifts, and force perturbations such as inertial, viscous, complex force fields), the generalization function can be narrow (10, 16, 21), broad (9, 16, 22–24) or non-monotonic (6, 10, 19). Even within a particular task, the pattern of generalization can differ depending on the variability of the training set (16), level of attentional distraction (25), the statistics of the perturbations(10, 26) and the visual workspace context (27, 28). Why is so much heterogeneity observed?

The heterogeneity in motor generalization functions seems unlikely to be easily explained by a model based solely on joints, muscles, and neural tuning basis sets. Ignored in most conceptions of generalization is how top-down inferences about how one should move in a new context may also influence motor learning (1), and, as such, influence generalization. This is surprising given that explicit, top-down movement strategies have been shown to be a significant component of behavior across a wide range of motor adaptation tasks (20, 29–32), and even to affect generalization directly (33–36).

In our view, some insufficiencies in our understanding of movement generalization can be amended if typical motor generalization tasks are, at least in part, conceptualized as normative decision-making problems (1, 4): How should one move in a new situation? In this perspective, generalization behavior can be thought of as including both a top-down decision-making component, and a bottom-up kinematically determined component; that is, motor generalization functions should be multiplexed, stemming from representations in both “psychological space” (3) and physical/kinematic neural coding space (11). The idea that generalization behavior is affected by distances between contexts in psychological space has unified seemingly disparate generalization behaviors across many task domains (3, 37–39). Thus it should be possible to observe motor generalization over both kinematic and psychological dimensions.

To test this, we performed five experiments that measured people’s explicit subjective judgments about different movement contexts, and their generalization of learned motor behaviors across those contexts. We found that the generalization of learned sensorimotor transformations includes a component that is strongly correlated with subjective similarity judgments about different movement contexts. Specifically, motor generalization can be affected by the similarity between training and transfer contexts in both physical (e.g, movement direction) and psychological (e.g., target shape) dimensions based on which of those dimensions is deemed relevant by the subject. This effect appears to map onto top-down decision-making processes in motor learning (40). Alternatively, implicit motor adaptation generalization appears to be shaped by physical dimensions. These findings broaden the range effects that top-down strategies have on motor learning, and also suggest that unless implicit motor learning is fully isolated, the generalization of adapted motor behavior will reflect multiple qualitatively distinct representational formats.

## Results

### Experiment 1

Here we asked if transfer of participants’ adapted motor behaviors from a single training target direction to novel target directions (Figure 1A) is related to subjective similarity between those contexts. Participants performed a match-to-sample similarity judgment task (Figure 1B), using a typical likert scale of 1-7 (with 1 reflecting the largest difference between contexts) to make pairwise comparisons between the act of reaching to a single “anchor” training target versus 45 unique probe target directions (Figure 1A, B, C; see *Methods* for details). We then imposed a visuomotor rotation of the visual feedback cursor at the anchor target direction (Krakauer et al., 2000), requiring participants to adapt their movements to restore performance at that target (Figure 1D). Lastly, we tested generalization by again having participants move to the probe target directions with no feedback.

**Figure 1.**
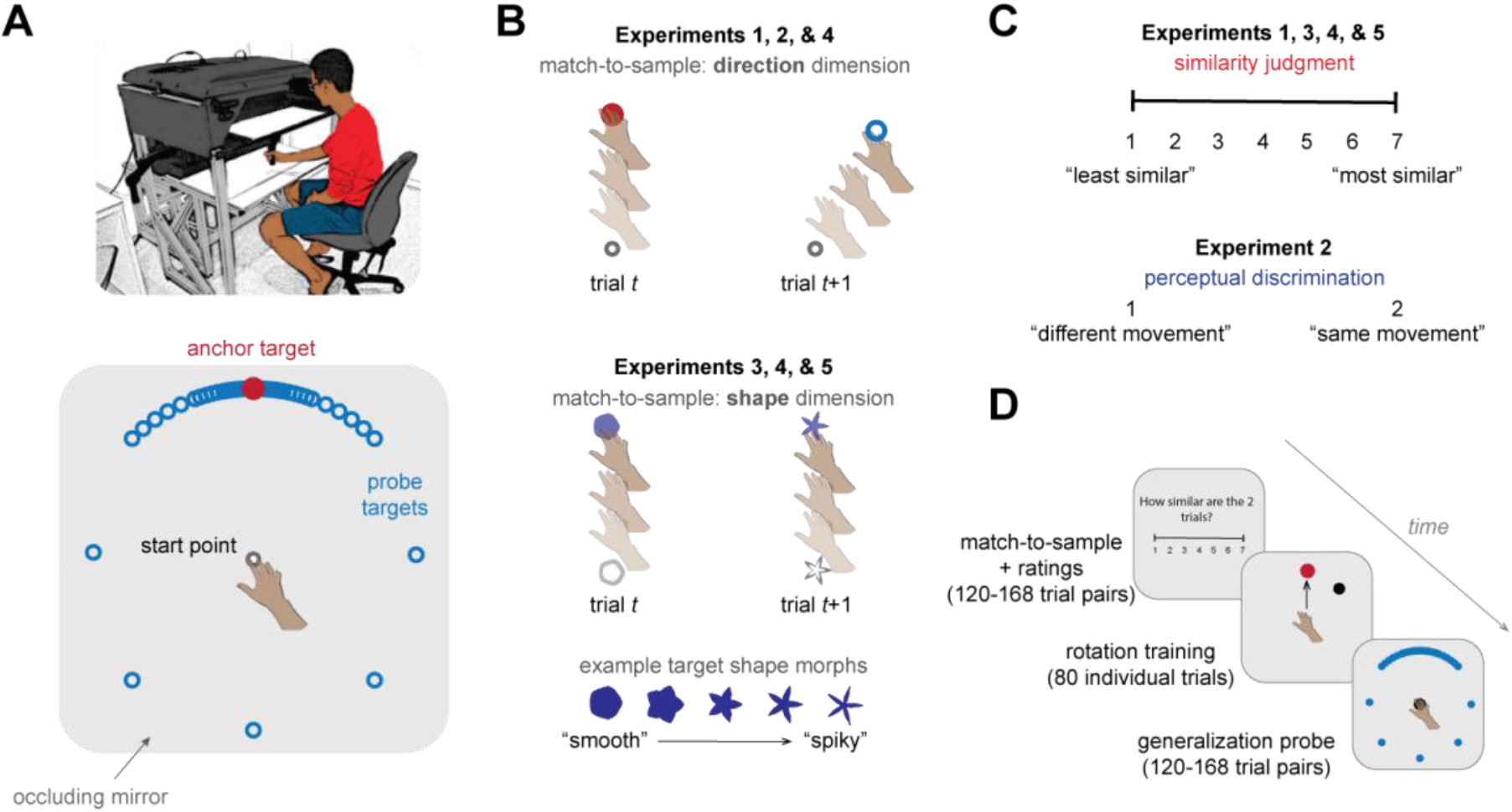
Experimental design and procedure. **(A)** Participants made center-out reaching movements using a robotic manipulandum (top). A schematic diagram of the task display (bottom) shows example target locations (blue circles) with respect to an anchor target (red). The hand and arm were occluded by a reflective mirror that displayed visual stimuli presented on a horizontally-mounted LCD screen. Only a single target is displayed per trial. **(B)** Match-to-sample judgment task. In Experiments 1, 2 and 4, participants performed trial pairs where they reached first to the designated anchor target location, then to a probe target in a different location. A similar method was used in Experiments 3, 4 and 5, but instead of probe targets being placed in a new location, they were in the same location as the anchor target but continuously varied from the anchor target in an abstract “shape dimension.” The shape dimension ranged from “smooth” to “spiky”, and was based on a previous study (van Dam & Ernst, 2015). **(C)** After performing each match-to-sample trial pair, participants either rated the similarity between the two trials (Experiments 1, 3, 4, and 5) or their perceptual discriminability (Experiment 2). **(D)** Each experiment consisted of a rating phase, a rotation adaptation phase (45° rotation of visual endpoint feedback), and a generalization probe phase, in that order. The particular length of each phase varied across experiments (see *Methods*).

As illustrated in Figure 2A, participants quickly and robustly adapted to the 45° rotation imposed at the anchor target (asymptote: average of last 10 trials of learning, 42.85 ± 6.11°; mean ± 95% C.I.). During the generalization phase, participants’ movements to probe targets reflected the canonical motor generalization curve typically observed in adaptation tasks (e.g., (16)), with the degree of adaptation falling off exponentially with the distance of the probe from the training direction (Figure 2B). Participants’ generalization curves in the similarity judgment task showed a qualitatively similar shape, with decreasing similarity at further distances from the anchor/training target (Figure 2C, D).

**Figure 2.**
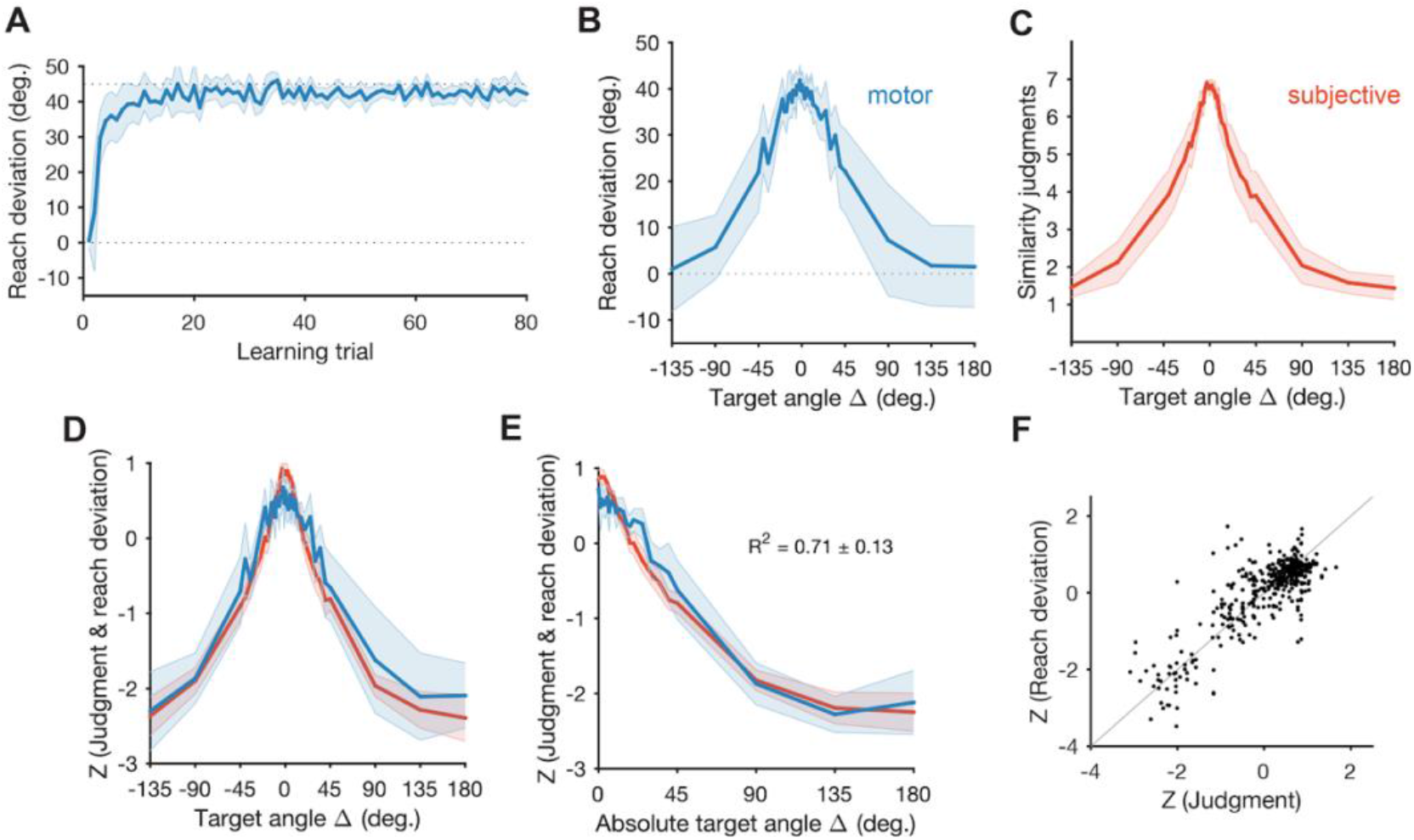
Experiment 1: Motor generalization reflects subjective similarity of different movement contexts. **(A)** Adaptation curve, reflecting the time course of participants learning to counteract the 45° rotation imposed at the anchor target. **(B)** Motor generalization function after adaptation to the rotation. The abscissa reflects the angular distance of the probe target from the anchor target (with the anchor target fixed at 0° for all participants for visualization purposes), and the ordinate reflects the reach deviation with respect to a direct reach to the probe (i.e., 0° deviation). **(C)** Subjective similarity function, participants performed a match-to-sample task, reaching to an anchor target direction and then a probe target direction, and then reporting on a 1-7 scale the similarity between the two movements (1 = least similar; 7 = most similar). **(D)** Normalized (z-scored) data from panels (C) and (D). **(E)** Decay functions were computed after normalization by collapsing target direction based on absolute angular distance. **(F)** Visualization of pooled data consisting of each participant’s normalized generalization/rating data (gray line = identity line). Error shading = 95% C.I.

The key analysis in Experiment 1 involved directly comparing the gradients of the subjective and motor generalization functions. We folded each generalization function using the absolute angular distance of probe targets from the training (anchor) target and z-scored both functions (within each condition and subject) to afford direct comparisons (Figure 2E). Each function showed roughly monotonic exponential decay, consistent with Shepard’s Universal Law of Generalization (3). We used linear regression to statistically compare each participant’s average similarity and motor generalization functions. Consistent with our hypothesis, we observed a striking agreement between these functions (Figure 2E, F): the similarity judgment curve provided a strong fit to the motor generalization curve, with an *R^2^* value of 0.71 ± 0.13. Critically, while both functions were expected to monotonically decrease with greater probe distance, there was no *a priori* reason, apart from our hypothesis, to believe that they would decay at nearly equivalent rates. Experiment 1 thus suggests that motor generalization may reflect in part a “readout” of subjective inferences about the relationship between learning and transfer contexts (i.e., target locations).

### Experiment 2

Experiment 2 served two functions: First, this experiment tested the alternative hypothesis that perceptual confusability between target directions better explains motor generalization; second, this experiment acted as a control for a potential confounding factor in Experiment 1, wherein the act of making target direction similarity judgments may have biased motor behavior. The procedures for Experiment 2 were identical to Experiment 1, but instead of making similarity judgments participants simply reported whether they thought two different reaches, directed at the anchor target and a probe target, were of the same or different direction (Figure 1C).

Participants in Experiment 2 also adapted easily to the perturbation (asymptote: 42.38 ± 4.63°; Figure 3A). Motor generalization in this new sample mirrored that seen in Experiment 1, showing a wide Gaussian function and gradual decay with increasing probe distance (Figure 3B). In contrast, perceptual confusability between the anchor and probe targets rapidly disappeared as the angular difference increased, with a just noticeable difference (50% JND) of 4.80 ± 0.71° (Figure 3C; JND determined via exponential fits; see *Methods*). As seen in Figure 3D, perceptual report functions captured a modest amount of variance in motor generalization (*R^2^* = 0.20 ± 0.06).

**Figure 3.**
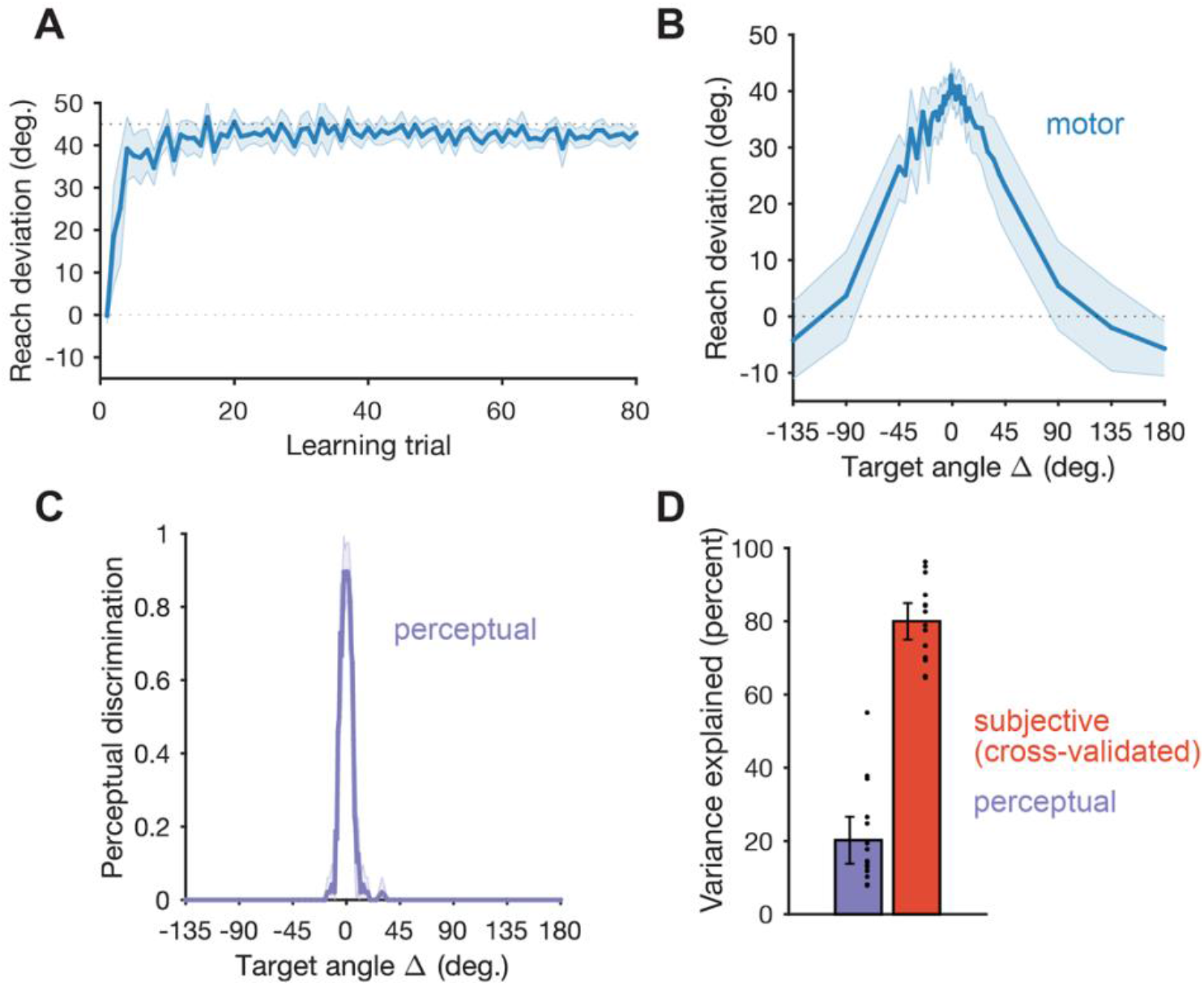
Experiment 2: Motor generalization is better explained by subjective ratings of contextual similarity versus perceptual discriminability. **(A)** Adaptation learning curve. **(B)** Motor generalization function for Experiment 2. **(C)** Average perceptual discriminability of reaches to the anchor target versus probe targets at various angular distances (a value of 1 reflects reaches judged as identical; a value of 0 reflects reaches judged as different). **(D)** Motor generalization variance explained by participant’s perceptual discrimination functions (violet) versus the average similarity rating function measured in the separate group of participants performing Experiment 1 (red). Dots represent individual participants. Error shading and error bars = 95% C.I.

A cross-validated analysis pointed to a role for psychological similarity in motor generalization: We took the average similarity judgment decay function from Experiment 1 (Figure 2E, red) and regressed it onto the (folded) motor generalization function from Experiment 2. This out-of-sample similarity function explained 80% of the variance (*R^2^* = 0.80 ± 0.06) in the Experiment 2 motor generalization function (Figure 3D), reflecting a four-fold increase in variance explained when using similarity judgments from a separate sample versus participants’ own perceptual judgments (comparison of *R^2^* values, *t*(15) = 23.41; *p* < 0.001). In addition to replicating the findings from Experiment 1, these results also rule out a confound wherein the act of making similarity judgment biases motor generalization.

### Experiment 3

So far, our results have shown that explicit similarity judgments about reaching to different spatial locations correlates with the generalization function seen in adapted movements. Next, we asked if motor generalization functions match subjective judgments only when both are linked to a physical dimension (e.g., reaching direction), or any contextual dimension could influence motor generalization. To that end, in Experiment 3 we cued different contexts using a purely visual cue – target shape – rather than target direction (Figure 1B). Specifically, the target’s shape on each trial reflected a single “morph” in an equispaced linear mapping from “round” to “spiky” shapes (41). For all trials and participants, the target was always presented at a fixed location regardless of its shape. Apart from these changes, the experiment mirrored Experiment 1 (save for minor alterations to the length of each task phase; see *Methods*). Participants judged the similarity between their particular anchor shape (i.e., either the most rounded or most pointed shape, counterbalanced across participants) and each of 23 probe morphs. They then adapted to a 45° visuomotor rotation imposed on the anchor training shape, and subsequently generalized to novel probe shapes.

Motor adaptation was again rapid and robust, as observed in the previous experiments (Figure 4A; asymptote: 43.12 ± 6.17°). Critically, participants showed monotonically decreasing motor generalization as a function of the target morph distance (in shapespace) from the anchor training target (Figure 4B). Again, we observed a similar decay function for the similarity judgments (Figure 4C, D; *R^2^* = 0.85 ± 0.04). We emphasize here that the similarity between these functions is parametric and suggests a subtle behavioral process, where during the generalization phase, participants directed their movements to angles intermediate between the target and rotation solution (e.g., 20°) when the target shape itself was similarly intermediate. We note that the motor generalization functions for direction (Figures 2 and 3) versus shape (Figure 4) have slightly different shapes, an observation we address in the next experiment. Overall, these data suggest that motor generalization can, in part, be affected by the distance between training and transfer contexts in both physical and abstract psychological dimensions.

These data so far are broadly consistent with the Universal Law of Generalization, which describes that an abstract representational space shapes generalization behavior across a diverse range of tasks(3). However, a key ingredient from that proposal is missing here: The dimension over which generalization occurs must be deemed “consequential” by the learner or it will be ignored. That is, if a butterfly’s color correlates with it being poisonous but its wing size doesn’t, a bird should generalize with respect to the former whilst ignoring the latter (assuming they are orthogonal). So far, our tasks have only had a single salient dimension (i.e., direction or shape). It is perhaps unsurprising then that participants inferred that this salient dimension is relevant to the learning task, and thus parametrically generalized over it. In Experiment 4 we pitted these two dimensions against each other and biased participants through different explicit instructions to deem one dimension as “relevant” and ignore the other. If generalization is indeed shaped by the so-called consequential dimension, we expect participants’ motor behavior to generalize according to the dimension emphasized by our instructions.

### Experiment 4

Two separate groups of participants were exposed to an identical sequence of trials, starting with a similarity judgment phase, an adaptation phase, and a generalization probe (such as illustrated in Figure 1D). Target location and target shape morph were simultaneously varied in a fully factorized design (i.e., an equal number of unique target morphs were experienced at each target direction). Crucially, participant group was determined by the instructions they received from the experimenter: In the Direction Emphasis group, participants performed direction-based similarity judgments and were told to attend to target direction and “adjust based on how similar the movement directions are”; in the Shape Emphasis group, participants performed shape-based similarity judgments and were told to attend to target shape and “adjust based on how similar the shapes are”. (We note again that we purposefully did not specify what type of adjustment participants should make; see *Methods*).

As depicted in Figures 5A and 5B, participants in both groups readily adapted to the rotation (asymptotes: 43.18 ± 5.37° and 42.07 ± 4.6°, respectively). Instruction had a strong effect on generalization: When looking at the effect of target shape (Figure 5C), the Shape Emphasis group showed the predicted motor generalization function, replicating the results of Experiment 3. Conversely, the Direction Emphasis group did not generalize according to shape. That is, because each shape was seen an equal number of times in each direction, the Direction Emphasis group’s shape-based generalization function was a nearly flat line centered on their average movement angle during the probe trials.

**Figure 4.**
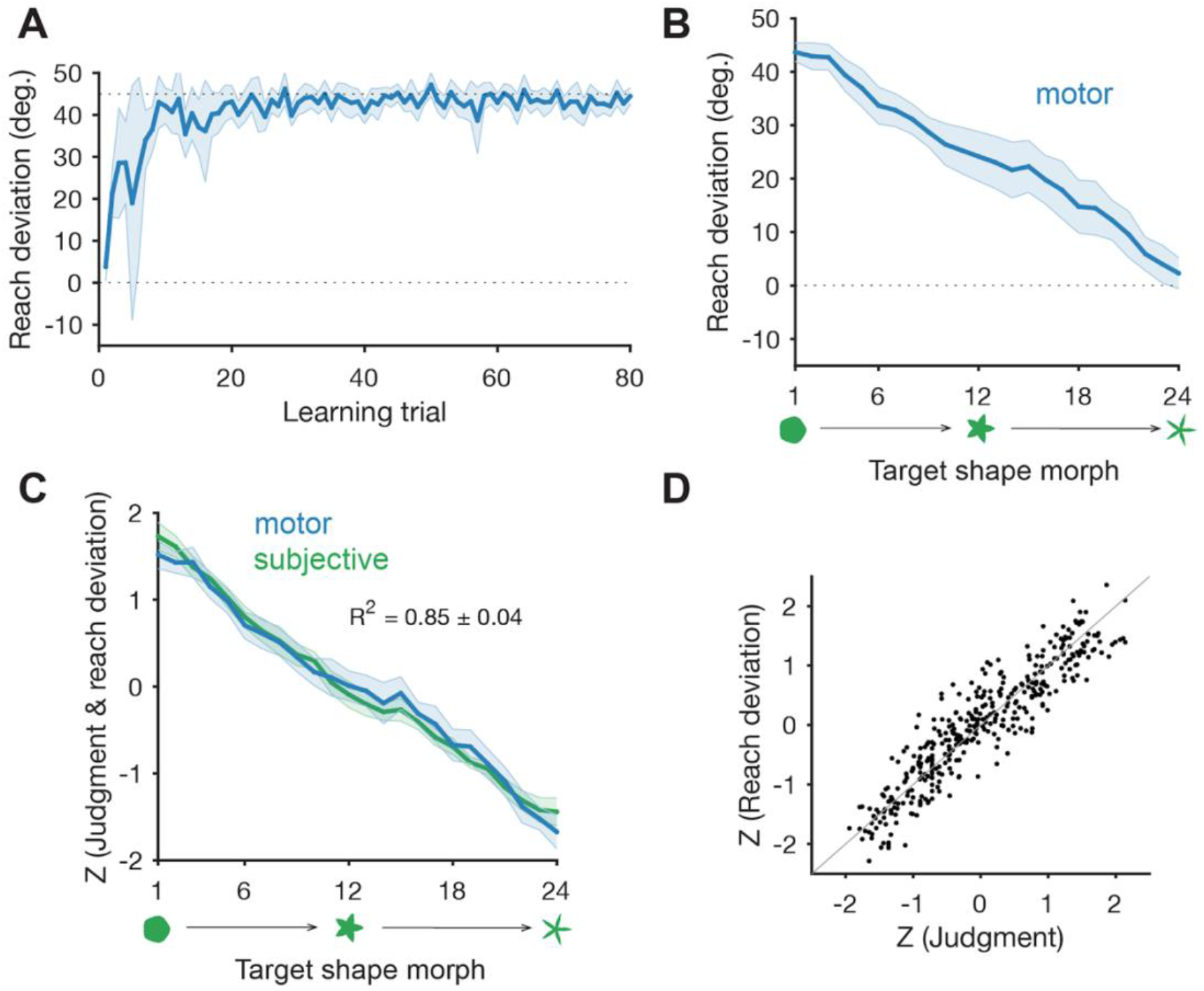
Experiment 3: Motor generalization is shaped by an abstract psychological space. **(A)** Adaptation curve. **(B)** Motor generalization as a function of target shape (target direction was held constant in all trials). **(C)** Normalized motor generalization (blue) and similarity judgment (green) functions, with respect to target shape. **(D)** Visualization of each participant’s normalized generalization/rating data (grey line = identity line). Error shading = 95% C.I.

**Figure 5.**
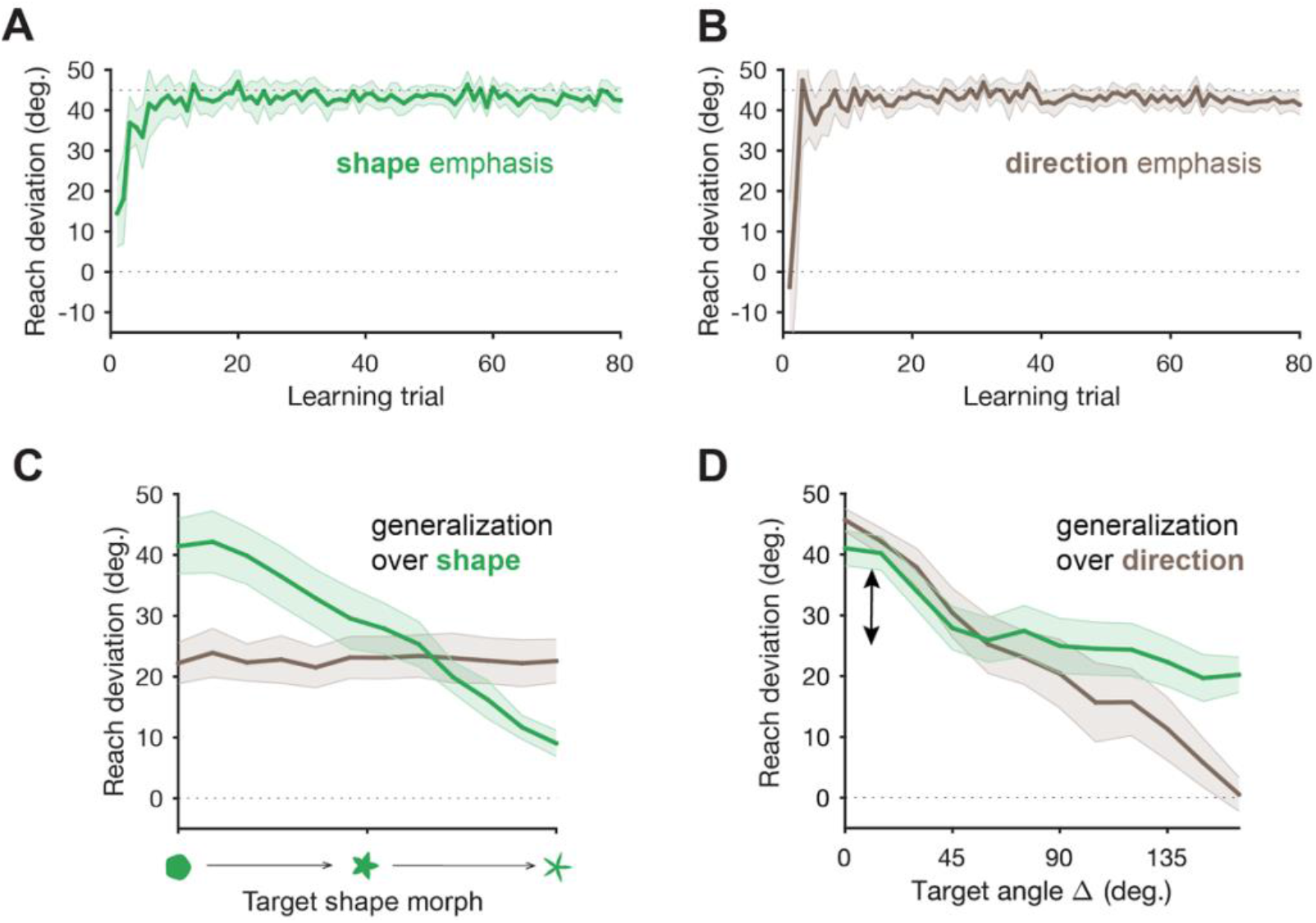
Experiment 4: Generalization is influenced by explicitly emphasized contextual dimensions. Adaptation curves in the Shape Emphasis **(A)** and Direction Emphasis **(B)** groups. **(C)** Motor generalization curves with respect to target shape, for the Shape Emphasis group (green) and Direction Emphasis group (brown). **(D)** Motor generalization curves with respect to target direction, for the Shape Emphasis group (green) and Direction Emphasis group (brown). The black double arrow highlights generalization behavior in the Shape Emphasis group (green) over the “irrelevant” dimension of direction. Error shading = 95% C.I.

When generalization was quantified with respect to target direction (Figure 5D), we found similar results, though we also observed directional biases even when the direction dimension was instructed to be non-consequential. That is, while the Direction Emphasis group showed the predicted generalization function as target distance increased from the training target, the Shape Emphasis group’s generalization behavior was also affected by target direction: In the Shape Emphasis group, generalization near the training target location was 17.36 ± 4.00° higher than the mean reaching deviations beyond the 45° probe target (*t*(15) = 8.51; *p* < 0.001). These results support two conclusions: First, that motor generalization is influenced by the explicitly consequential task dimension, and second that movement direction influences motor generalization even when it is not supposed to be relevant.

The observed obligatory effect of movement direction has a likely source – motor adaptation has a prominent implicit learning component, and this component is thought to generalize in these tasks according to kinematic dimensions (14, 16, 20, 42). Thus, we hypothesized that generalization over the irrelevant directional dimension seen in Figure 5D (green) was likely due to an implicit component of motor learning. To test this, in our next experiment we isolated explicit and implicit components of visuomotor adaptation, predicting that the generalization of explicitly learned visuomotor transformations would scale with subjective similarity in psychological space. In contrast, the generalization of implicitly learned visuomotor transformations would not be significantly modulated by non-kinematic dimensions (i.e., target shape). This observation would suggest that motor generalization is influenced by a mixture of factors, including top-down inferences and kinematically-linked implicit representations.

### Experiment 5

We isolated putative explicit and implicit processes of motor learning by using delayed endpoint feedback (43) and task-irrelevant error-clamp feedback (42) respectively. Delayed endpoint feedback has been shown to selectively disrupt the process of implicit adaptation and thus appears to isolate predominantly explicit forms of learning during a visuomotor rotation task (43). In contrast, for clamped visual feedback, a visuomotor rotation is imposed such that a visual feedback cursor is yoked to the participant’s reaching velocity but follows a rigid path at a fixed deviation from the target (see *Methods*); even though participants are instructed to ignore this feedback, their movements incrementally deviate in the direction opposite the error without their awareness (44)

We replicated the procedure in Experiment 3 (i.e., shape-based generalization) using delayed feedback for the Delay group and clamped error feedback for the Clamp group, as outlined above. As illustrated in Figure 6A, while both groups displayed a rapid change in hand angle in response to the visuomotor rotation at the anchor training shape, they displayed considerably different adaptations to the rotation. This indicates the expected qualitatively different forms of learning when using delayed endpoint feedback and error-clamp feedback during visuomotor rotation learning. Participants in the Delay group explicitly re-aimed near the 45° solution under delayed endpoint feedback imposed at the anchor training shape, reaching an asymptote of 45.24 ± 6.92° (averaged over the last 10 trials of training). In contrast, participants in the Clamp group implicitly adapted 8.68 ± 3.45° to the 20° error-clamp imposed at the anchor training shape.

**Figure 6.**
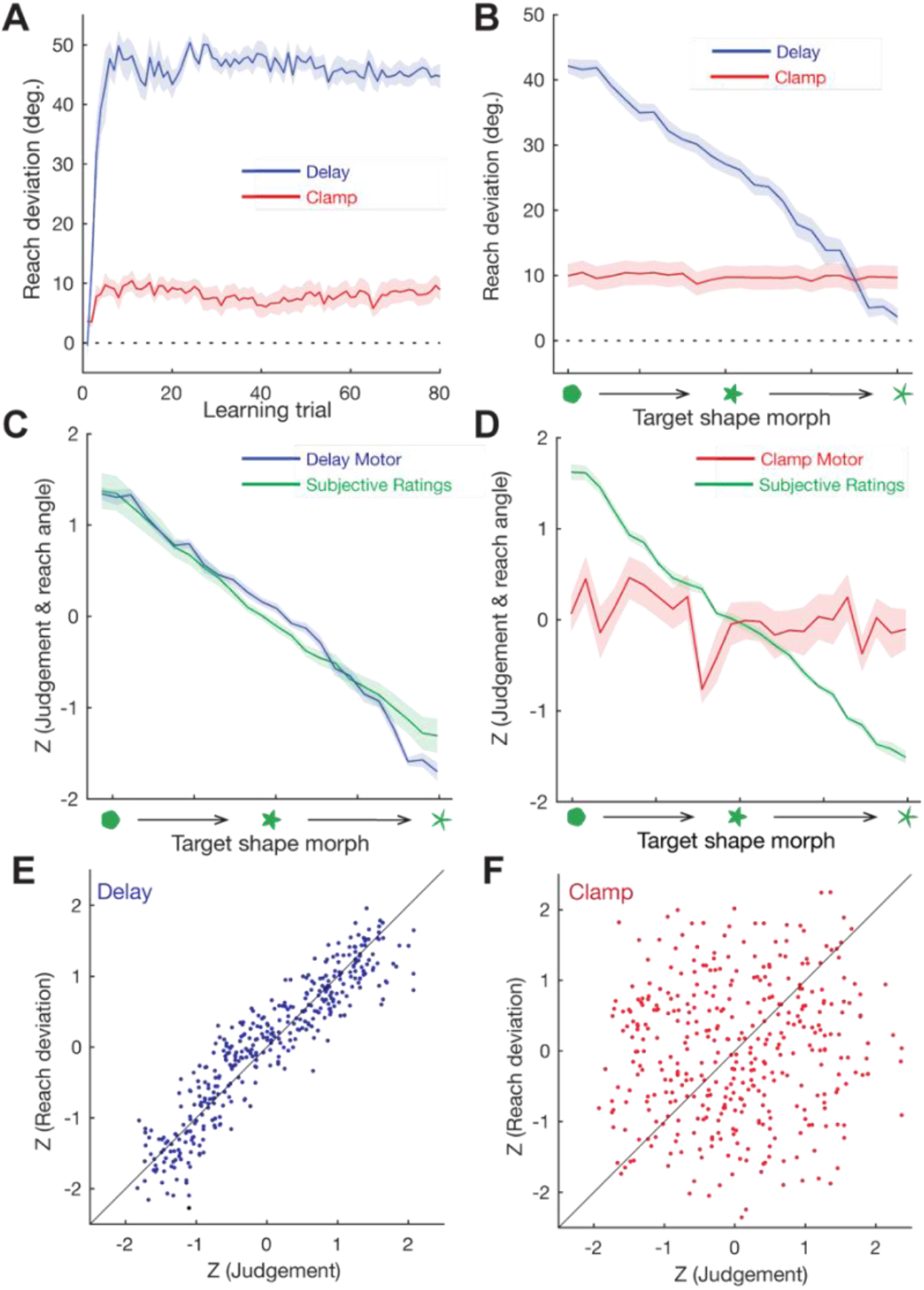
*Experiment 5:* Isolated explicit and implicit processes of learning *were differentially modulated by subjective inference*. **(A)** Participants learned to compensate for a visuomotor rotation using delayed endpoint feedback (blue) and task-irrelevant error-clamp feedback (red). They displayed considerably different adaptations to the rotation. **(B)** Targets were only presented in a single direction, but with 24 different shape morphs, with the anchor target on the extremes of the target shape dimension. Explicit generalization monotonically decreases as a function of the shape morphs. In contrast, implicit adaptation generalized similarly across shape morphs. **(C)** Normalized explicit motor generalization (blue) and subjective similarity ratings of target shapes (green) were significantly related. **(D)** Normalized implicit motor generalization (blue) and subjective similarity ratings of target shapes (green) were not significantly related. **(E)** and **(F)** Visualization of each participant’s normalized generalization/rating data (gray line = identity line) for explicit (E) and implicit (F) components of learning. Error shading = 95% C.I.

Generalization of explicitly and implicitly learned visuomotor transformations was differentially modulated by non-kinematic dimensions (i.e., target shape): Explicit generalization monotonically decreased as a function of the target shape morphs (Figure 6B). In contrast, implicit generalization across target shapes was strikingly uniform – participants moved in roughly the same (adapted) direction regardless of the shape of the probe target (Figure 6B; mean slope of generalization function: −0.01 ± 0.01); we note that while small, there was a marginal negative effect of probe target shape dissimilarity versus the anchor on motor generalization (*t*-test on regression slopes: *t*(15) = 1.93; *p* = 0.07).

Subjective similarity judgments replicated what was observed in Experiment 3, showing monotonically decreasing similarity ratings as the probe morph deviated from the anchor target shape for both the Delay (explicit) and Clamp (implicit) conditions. We observed a strong correlation between similarity judgements and explicit generalization functions (Figure 6C), with an *R^2^* value of 0.85 ± 0.03. However, subjective similarity between target shapes appeared to play a negligible role in implicit motor generalization (*R^2^* = 0.06 ± 0.02). These findings suggest that generalization of motor behavior in psychological space probably maps onto a top-down intentional component of motor learning (40), whereas obligatory directional generalization is primarily the consequence of implicit learning (20, 29, 33, 34, 42).

## Discussion

Shepard’s “Universal Law of Generalization” (3) states that the probability of a learned behavior (e.g., a pigeon’s knowledge that pecking at a blue light flash predicts a reward) generalizing to a novel context (e.g., a green light flash) falls off exponentially with the putative “psychological distance” between contexts (e.g., subjective similarity) rather than their physical distance (e.g., the wavelength of light). The mental representations that maintain these so-called psychological distances supposedly determine how an agent infers an adaptive course of action in similar (or dissimilar) situations. Psychological distance is thought to encompass the full space of arbitrary task and stimulus dimensions, and is typically quantified using methods like multidimensional scaling (45). This approach to understanding generalization has successfully unified a wide variety of seemingly disparate generalization behaviors observed in humans and animals (37, 46).

In the current experiments, we tested the straightforward idea that subjective similarity can explain some aspects of motor generalization. To do this, we measured people’s subjective similarity judgments over both a physical dimension believed to dictate motor generalization (movement direction) and a novel abstract dimension (target shape). We found that subjective judgments about different physical contexts could explain a significant portion of variance in the generalization of learned sensorimotor behaviors (Experiment 1). We also found that the correlation between subjective similarity and motor generalization was above and beyond what could be explained by perceptual constraints (Experiment 2) or kinematics alone (Experiment 3). Explicit instructions about the “consequential” (task-relevant) dimension of different movement contexts determined how people generalized (Experiment 4). Lastly, we confirmed that the above effects were likely all related to top-down, intentional aspects of motor learning rather than implicit motor adaptation (Experiment 5). Our five experiments support a key role for cognitive judgments in shaping motor generalization. The underlying factor informing these judgments could be described as the psychological distance between training and transfer contexts.

In contrast to a broad framework like the concept of “psychological distance,” the framework typically used to describe generalization of learned motor behaviors is usually more domain-specific – generalization is thought to be driven exclusively by movement-specific tuning constraints in motor regions of the brain (6, 8–10, 13–18) (e.g., primary motor cortex, cerebellum, etc). A unifying perspective on the heterogeneous generalization behavior seen in human motor learning studies has been elusive. We suggest that this is partly because the scope of representations influencing motor generalization has focused primarily on movement directions, postures, joint positions, and the like. People’s explicit judgment about which action they ought to take in a given context should be considered as another key component of motor generalization.

There is a growing body of work showing that many aspects of human motor learning can be traced to top-down strategies that operate alongside implicit processes (29, 32, 40, 47). Recent work has shown that performing even relatively simple motor learning tasks is a cognitive endeavor, leveraging planning and mental imagery (48, 49), working memory (30, 50, 51), and long-term memory (52–54). Interestingly, these top-down factors do appear to directly affect motor generalization, even when generalization is linked to implicit learning – when participants are asked to report their intentions during adaptation tasks (i.e., verbalize where they are aiming their movements), explicit verbal reports capture much of the observed generalization function versus implicit adaptation alone (29, 34, 36) and these cognitive strategies appear to directly influence the dynamics of implicit learning (33–36, 49). Future research could target the role of subjective similarity in implicit learning in situations that allow both explicit and implicit learning to interact.

Our study has several limitations. First, we exclusively used explicit reports (subjective similarity ratings) to characterize psychological distances between movement contexts. While Experiment 2 controlled for the effect of making subjective similarity judgments on motor generalization, future studies could use more subtle methods to estimate an individual’s contextual judgments, such as having them perform secondary decisionmaking tasks with the relevant stimuli (e.g., classification tasks). We deliberately chose not to record explicit aiming reports to quantify cognitive aiming strategies, as has been done in previous work on visuomotor learning (47). While these data can be informative, we decided against this method primarily to avoid interfering with (or priming) participants’ volitional strategies, which is a potential consequence of requiring them to report their intentions (1, 55, 56).

Second, visuomotor adaptation is a relatively simple motor learning task. In the future, more challenging, ecologically relevant motor learning tasks (57) could be used to examine the role of top-down inferences in real-world skill learning. For example, how do high-level inferences about novel tools shape how we initially decide to physically interact with them? We would expect similar results across a range of skill learning tasks.

Third, our methods involved directly measuring participants’ contextual judgments, eschewing the need to quantitatively model the inference process. How should we model inference in motor generalization, particularly when we do not have direct access to participants’ judgments about different situations? One promising approach would be to leverage Bayesian models of generalization from more cognitive domains (4) and apply them to motor tasks. Indeed, probabilistic approaches have begun to be successfully applied to the learning of different sensorimotor policies during motor adaptation (1, 58). Clustering algorithms are another promising approach, such as those used in modeling the learning of context-specific action policies during reinforcement learning (2). Future studies using these computational techniques could be combined with neurophysiological data to characterize and anatomically locate the cognitive representations that determine the generalization of motor skills.

## Materials and Methods

### Participants

A total of 112 right-handed (Oldfield 1971) participants (67 females; age range: 18-36) were recruited from the research participation pool of the Department of Psychology at Princeton University for course credits or cash. All sample sizes were decided *a priori* and are similar to those in previous publications (14, 20), and also supported our counterbalancing requirements. Each participant was randomly assigned to one of the five experiments. All experimental protocols were approved by the Institutional Review Board at Princeton University.

### Apparatus and Reaching Task

The same apparatus was used in all five experiments in this study. Participants sat comfortably in a chair and made ballistic reaching movements while grasping the handle of a robotic manipulandum with their right hand (KINARM End-Point; BKIN Technologies). The participants’ arms were not provided with any external support, and all movements were restricted to the horizontal plane (see Figure 1A). All visual stimuli were projected to the participant via a horizontal display screen (LG47LD452C; LG Electronics) reflected onto a semi-silvered mirror mounted 6 cm above the robotic handle. The mirror occluded vision of the arm and hand (Figure 1A, B) and the robotic handle, preventing direct visual feedback of hand position. Movement kinematics were recorded at 1 kHz.

At the initiation of each trial, the robot rendered a spring-like load drawing the current position of the hand into a central starting location in the middle of the display. In Experiments 1 and 2, the central starting location was depicted by a gray empty circle 1 cm in diameter, while in Experiments 3-5, the starting location was represented by a gray hollow shape (diameter: ~1 cm) that varied from “smooth” to “spiky” (see description of target shapes below). When the hand was within the starting location, the gray starting location turned white and a cursor (white filled) corresponding to the shape of the starting location appeared. After maintaining the hand within the starting location for a random foreperiod (uniformly drawn from 100-1000 ms), a “beep” tone coincided with the presentation of a target (blue-filled shape) at a radial distance of 9 cm. Participants were then instructed to perform a ballistic reaching movement that “sliced” through the target.

Depending on the particular experiment and training conditions, movements were performed under one of four different forms of feedback: continuous cursor feedback, clamped feedback, delayed endpoint feedback or no feedback. On continuous feedback trials, feedback of the cursor remained visible throughout the duration of the outbound movement. Once movement extent exceeded 9 cm, the visual feedback of the cursor froze on the screen to provide static feedback of the final hand position for 1 s. On clamped feedback trials, the cursor was visible during the outbound movement and followed an invariant trajectory that was offset by ±20° relative to the target, regardless of the hand movement direction (42). Once the movement extent exceeded 9 cm, the cursor was frozen on the screen for 1 s to provide static feedback of the final hand position when the workspace radius was crossed. On delayed endpoint feedback trials, visual feedback of the cursor position was withheld when movement velocity exceeded 5 cm/s. Endpoint feedback of the cursor subsequently appeared on the circumference of an invisible ring of radius 9 cm after 1.5 s (20, 43). Finally, on no feedback trials, the cursor was hidden throughout the trial.

### General Experimental Procedures

We performed five experiments that measured people’s subjective judgments about different movement contexts, and their generalization of learned motor behaviors across those contexts. All experiments followed the same general experimental procedures which consisted of three phases. The first phase was a rating phase in which participants either judged which movement context presented was more similar to a reference movement context (Experiments 1, 3-5) or determined if they were perceptually similar to each other (Experiment 2). This was followed by a rotation adaptation phase in which all movements were performed under a visuomotor rotation confined to a single training/movement context (a single target direction/target shape). In the final generalization probe phase, we continued exposure to the visuomotor rotation in the training context while examining generalization of the learned motor behavior across different movement contexts using no feedback probe trials. The length of each phase varied slightly across experiments depending on the training conditions (see below).

### Design of Experiment 1: Movement direction subjective similarity and generalization

Experiment 1 (n=16) investigated if motor generalization from a single learning target direction to novel target directions (Figure 1A) echoed the subjective similarity between different movement direction contexts. The experiment started with a rating phase where participants made pairwise comparisons between the act of reaching to a single “anchor” training target direction (chosen from 8 possible locations: 45°, 90°, 135°, 180°, 225°, 270°, 315°; counterbalanced across participants) versus 45 unique probe target directions (Figure 1B). The 45 probe target directions spanned the range of ±180 with respect to the anchor target and were spaced at 1° intervals between 0 and ±10, 2° intervals between the range ±10 to ±20, 5° intervals between the range of ±20° to ±45°, and 45° intervals from ±45° to ±180°. This range was chosen so as to best capture the sensitivity of subjective ratings and the shape of the generalization functions observed in pilot experiments. The distribution of probe targets was also consistent with the range of movements examined in most previous studies of motor generalization. All movements were performed under continuous online feedback, with veridical feedback.

After each pair of movements, participants were asked to judge “How similar were the two trials?” and verbally reported similarity based on a likert scale of 1-7 (with 1 being extremely dissimilar and 7 reflecting extremely similar). The trial pairs were not speeded, and the likert scale remained visible on the display until the participants responded. It is important to note that participants were not instructed to pay particular attention to any differences in movement kinematics (i.e. direction, speed, amplitude), and instead were asked to render their judgments based on the perceived contextual similarity between the pair of trials. In all, participants made 138 judgments of subjective similarity between movement pairs (6 repeats for each unique anchor and probe movement pair).

Next, in the rotation adaptation phase, participants completed a visuomotor rotation training task which consisted of 80 individual movements with a ±45° rotation imposed on the visual feedback cursor (the sign of the rotation was counterbalanced across participants). All learning trials were performed at the anchor target direction. On a typical initial exposure to the visuomotor rotation, participants moved their hand movement directly toward the anchor target direction which resulted in a cursor movement that veered off course by 45° in relation to the center starting location, as shown in the middle panel of Figure 1D. This required participants to adapt their movements in the opposite direction of the applied visual rotation to restore performance at the anchor target direction (Figure 1D).

Finally, in the generalization probe phase, participants completed 138 pairs of movements to assess the transfer of learned movements from a single learning target direction to novel target directions. The trial structure was similar to that in the rating phase and only differed in the type of feedback presented during the pairs of movements. Specifically, for each pair of movements, continuous visual feedback was provided for movements to the anchor target direction to reinforce the visual rotation that was previously learned. All reaches to probe targets were no feedback trials, allowing us to measure generalization.

### Design of Experiment 2: Perceptual confusability control experiment

Experiment 2 (n=16) tested the alternative hypothesis that it is the perceptual confusability, and not subjective similarity, between learning and transfer contexts that influences motor generalization. Moreover, this experiment also controlled for any confounding effects that the act of making subjective similarity judgments may have on motor learning and generalization. The experimental procedure was similar to that of Experiment 1, except that participants were asked to judge if the two movement contexts presented were the *same* during the initial rating phase. That is, participants made pairwise comparisons between the act of reaching toward a single “anchor” training target versus the 45 unique probe target directions and were simply tasked with discriminating, i.e., “Were the 2 trials the same?” The question remained visible on the display until participants verbally responded yes/no and the experimenter recorded the response and initiated the next trial.

### Design of Experiment 3: Abstract contextual dimensions and motor generalization

So far we have focused largely on similarity judgements over traditional contextual dimensions, such as target direction, which is the dominant method for studying motor generalization. To determine if the relationship between similarity judgements and motor generalization can be applied to a more abstract contextual dimension, Experiment 3 (n=16) measured subjective similarity and motor generalization across different target shape “morphs.” These targets were always presented at a single fixed location.

The target shape morphs used in the experiments were chosen from an equispaced linear mapping from “round” to “spiky” shapes used in a previous study (van Dam and Ernst, 2015). Briefly, each target shape morph differs in the length of each of its five points (the outer radius ro) and in the length of the intervening indents (the inner radius ri; see Figure 1, van Dam & Ernst, 2015). Every shape in this “pointiness” scale can be described by a shape parameter *p*, which takes the ratio between the outer and inner radii:

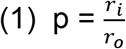

Here, a shape parameter of 1 indicates that outer and inner radii are equal and therefore corresponds to a typical round shape. In contrast, a small *p* indicates that the outer radius is much larger than the inner radius and the shape is perceived as “spiky”. In the current experiment, we selected 24 shapes where *p* was evenly spaced between 0.1 to 0.9 (See Figure 1B). (For detailed information about how each target shape morph was constructed see “Mathematical description of the target shapes” in (41))

During the rating phase, participants performed the match-to-sample subjective similarity judgment task (Figure 1B) to make pairwise comparisons between the act of reaching toward a single “anchor” training shape (either the most rounded or most pointed shape, counterbalanced across participants) versus 23 probe target shapes. We emphasize that, for all trials and participants, the target was always presented at a fixed location, regardless of its shape. After each pair of movements, participants judged, “How similar were the 2 target shapes” using a likert scale of 1-7 (with 1 being extremely dissimilar and 7 reflecting extremely similar). The likert scale remained continuously visible on the screen until participants verbally responded. In all, participants made 120 judgments of subjective similarity between the two target shapes (5 repeats for each unique anchor target shape and probe target shape pair).

In the rotation adaptation phase, participants completed 80 individual movements with a ±45° visuomotor rotation imposed on the visual feedback (rotation sign counterbalanced across participants), only being exposed to the anchor shape. Finally, the generalization probe phase consisted of 120 pairs of movements to assess generalization from the anchor training shape to novel probe shapes (5 repeats for each unique anchor target shape and probe target shape pair). Similarly, for each pair of movements, rotated visual feedback was provided on the first movements toward the anchor training shape, but feedback was not shown on movements toward the probe target shapes.

### Design of Experiment 4: Explicit contextual instructions and generalization

Experiment 4 (n=32) examined how instructions could bias participants toward different dimensions of the movement context (i.e., target locations versus target shapes), resulting in downstream effects on generalization. The experimental procedure was similar to Experiment 3, with two key differences: First, target location and target shape morph were simultaneously varied in a fully factorized design (i.e., an equal number of unique target morphs were experienced at each target direction). Second, slightly longer trial phases were used to account for the fully factorized design.

In the initial rating phase, participants compared movements made toward the anchor training target/shape (fixed at 0° or 180°, with the target shape either the most rounded or most pointed shape, counterbalanced across participants) and probe target movements which varied in direction (12 target locations spaced every 15° away from the anchor training target) and target shape (12 different shapes morphs from “round” to “spiky”). To emphasize particular contextual dimensions, two separate groups of participants were given different instructions: In the Direction Emphasis group, participants were told to attend to the target direction, and subjectively judge “How similar were the 2 movement directions?”, irrespective of target shape. In contrast, in the Shape Emphasis Group, participants were explicitly instructed to attend to the target shape and subjectively judge “How similar were the 2 target shapes?”, irrespective of the target direction. Overall, participants made 144 subjective similarity judgments (6 repeats for each unique anchor target location/shape and probe target location/shape pair).

The rotation adaptation phase consisted of 80 individual movement trials at the anchor training target location/shape with the visual feedback rotated by 45°. Finally in the generalization probe phase, participants performed a total of 144 pairs of movements to measure the generalization with respect to each contextual dimension. For the Direction Emphasis group, participants were instructed to “Adjust based on how similar the movement directions are” before each pair of movements, to probe generalization over the direction dimension. In contrast, in the Shape Emphasis group, participants were instructed to “Adjust based on how similar the shapes are” to probe generalization over the shape dimension. We note that no detail about the nature of this “adjustment” was given in the instructions, as to avoid biasing participants in any way.

### Design of Experiment 5: Subjective similarity and explicit/implicit motor generalization

Experiment 5 (n=32) investigated whether explicit or implicit processes underlie the generalization over abstract contextual dimensions. Moreover, this experiment was designed to further examine the small degree of generalization we saw in the Shape Emphasis group over the “irrelevant” dimension of target direction in Experiment 4 (Figure 5D), testing our hypothesis that this was due to the kinematic generalization of implicit learning. The experimental procedure mirrored Experiment 4, except that the type of rotated visual feedback provided during the rotation adaptation phase was different (see below), as were the length of the rating and generalization phases.

First, in the rating phase, participants performed a total of 168 pairwise comparisons to judge how similar the probe target shapes were to the anchor training shape (7 repeats for each unique anchor target shape and probe target shape pair). Next in the rotation adaptation phase, to measure explicit and implicit learning in relative isolation, participants were assigned either to a Delay condition (n=16) where we delayed their cursor endpoint feedback or a Clamp condition (n=16) where cursor feedback was clamped. Delayed endpoint feedback has been shown to selectively disrupt the development of implicit and appears to allow predominantly explicit forms of learning during a visuomotor rotation task (20, 43, 59). For this Delay condition, a 45° visuomotor rotation was introduced to a single anchor training direction/shape. In contrast, with clamped visual feedback, participants are firmly instructed to never aim their reaches elsewhere, aiming only directly toward the anchor training target and ignoring the offset feedback. This task has been shown to successfully isolate implicit learning (42); that is, participants’ movement directions undergo significant adaptation in the direction opposite of the error, and this learning occurs fully outside awareness (60). For this Clamp condition, the visuomotor rotation was imposed by restricting the path of the cursor along a constant 20° angle with respect to the target (with rotation direction counterbalanced across participants). We chose not to use a larger rotation size because pilot testing revealed that rotations larger than 45° did not produce a robust change in reach direction. Participants trained for a total of 80 trials under the delayed endpoint feedback or error-clamped visual feedback, before generalization was assessed to the probe target shapes in the generalization probe phase. Here, participants performed 168 movement pairs to assess the generalization of explicit and implicit motor learning over an abstract, non-kinematic dimension (target shape).

### Data Analysis

First, baseline movement biases were defined as the mean movement endpoint at each target location (or target shape) in the non-feedback match-to-sample/judgment block (Figure 1D). Movement endpoints were computed as the direction of the vector connecting the participant’s hand location at movement onset to the location when movements exceeded a radial distance of 9 cm. Movement onset was defined when the movement speed threshold first exceeded 5 cm/s. For each participant’s movement data, outliers were removed prior to analysis; we defined outlier movements as any movements that were more than 3 standard deviations from the mean movement direction across the experiment. This resulted in excluding, across the five experiments, an average of 0.18 ± 0.22% of trials (mean ± 95% C.I.). Generalization functions were quantified by taking the mean endpoint error at each probe location, and subtracting the relevant baseline bias. Adaptation learning curves for each experiment (e.g., Figure 2A) were similarly baseline-corrected and then averaged across participants. Generalization “decay” functions for experiments that utilized different target directions (e.g., Figure 2E) were computed by averaging data at each probe, with probes defined by their absolute angular distance from the anchor target. For visualization of generalization decay functions, we independently z-scored each average function (both rating and movement) for each participant.

Direct comparisons between generalization decay functions for subjective ratings versus motor behavior were conducted via linear regression, regressing the rating function of interest (e.g., similarity, perceptual) onto the movement generalization function. We then computed an R-squared metric for each participant to quantify the amount of movement generalization variance that could be explained by subjective judgments. Because we did not have similarity ratings in Experiment 2, we performed this analysis in a cross-validated manner, taking the mean similarity judgment function from Experiment 1 and regressing it onto each individual participant’s movement generalization function in Experiment 2. For all linear regression analysis, we verified the multivariate normality assumption of the data using a Kolmogorov-Smirnov test.

For the correlation analysis (Figure 7) we derived a decay rate from each generalization function. To do this, we fit their decay functions with an exponential function using the MATLAB *fit* function, with free parameters for a scaling factor (*a*) and a decay rate (*b*):

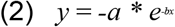

For between-participant correlations, we normalized (z-scored) decay rates for participants’ rating and movement generalization functions independently within each experiment, then computed the (Pearson) correlation between subjective and motor generalization functions pooling across experiments. This correlation analysis was limited to Experiments 1, 3, 4, and 5 as Experiment 2 did not have subjective ratings (Experiment 2). Similarly, to compute the 50% just-noticeable-difference (JND) in the perceptual discrimination task (Experiment 2), we fit an exponential decay function (Equation 2) to each participant’s perceptual reports (collapsed by absolute probe distance) and solved the resulting equation analytically at *y* = 0.50.

## Acknowledgments

We thank Carlo Campagnoli, Chandra Greenberg, and Adhvik Kanagala for help with study design and thoughtful discussions. This work was supported by the National Institute of Health under fellowship F32MH119797 awarded to S.D.M., and the National Science Foundation under grant number 1838462 awarded to J.A.T.

